# Structure of HIV-1 gp41 with its membrane anchors targeted by neutralizing antibodies

**DOI:** 10.1101/2020.11.12.379396

**Authors:** Christophe Caillat, Delphine Guilligay, Johana Torralba, Nikolas Friedrich, Jose L. Nieva, Alexandra Trkola, Christophe Chipot, François Dehez, Winfried Weissenhorn

## Abstract

The HIV-1 gp120/gp41 trimer undergoes a series of conformational changes in order to catalyze gp41-induced fusion of viral and cellular membranes. Here, we present the crystal structure of gp41 locked in a fusion intermediate state by an MPER-specific neutralizing antibody. The structure illustrates the conformational plasticity of the six membrane anchors arranged asymmetrically with the fusion peptides and the transmembrane regions pointing into different directions. Hinge regions located adjacent to the fusion peptide and the transmembrane region facilitate the conformational flexibility that allows high affinity binding of broadly neutralizing anti-MPER antibodies. Molecular dynamics simulation of the MPER Ab-induced gp41 conformation reveals the transition into the final post-fusion conformation with the central fusion peptides forming a hydrophobic core with flanking transmembrane regions. This, thus, suggests that MPER-specific broadly neutralizing antibodies can block final steps of refolding of the fusion peptide and the transmembrane region, which is required for completing membrane fusion.

## Introduction

Viral fusion proteins catalyze virus entry by fusing the viral membrane with cellular membranes of the host cell, thereby establishing infection. The HIV-1 envelope glycoprotein (Env) is a prototypic class I fusion protein that shares common pathways in membrane fusion with class II and III viral membrane fusion proteins ^1–4^. HIV-1 Env is expressed as a gp160 precursor glycoprotein that is cleaved into the fusion protein subunit gp41 and the receptor binding subunit gp120 by host furin-like proteases. Gp41 anchors Env to the membrane and associates non-covalently with gp120, thereby forming a stable trimer of heterodimers, the metastable Env prefusion conformation ^5,6^. Orchestration of a series of conformational changes transforms energy-rich prefusion Env into the low-energy, highly stable gp41 post-fusion conformation, which provides the free energy to overcome the kinetic barriers associated with bringing two opposing membranes into close enough contact to facilitate membrane fusion ^2,3^.

HIV-1 gp41 is composed of several functional segments that have been shown or suggested to extensively refold upon fusion activation: the N-terminal fusion peptide (FP), a fusion peptide proximal region (FPPR), the heptad repeat region 1 (HR1), a loop region followed by HR2, the membrane proximal external region (MPER), the transmembrane region (TMR), and a cytoplasmic domain. Structures of native Env trimers in complex with different broadly neutralizing antibodies revealed the conformation of the gp41 ectodomain lacking MPER in the native prefusion conformation ^7–12^. Env interaction with CD4 results in opening of the closed prefusion trimer ^13,14^, which includes the displacement of gp120 variable regions 1 and 2 (V1-V2) at the apex of the trimer but no changes in gp41 ^15^. This is required for the formation of a stable ternary complex of Env-CD4 with the co-receptor ^16–18^. Co-receptor binding positions prefusion gp41 closer to the host-cell membrane ^5^ and induces a cascade of conformational changes in gp41. First, the fusion peptide is repositioned by ~70 Å ^9^ to interact with the target cell membrane, generating a 110 Å extended fusion-intermediate conformation ^19,20^ that bridges the viral and the host cell membrane ^21^. Subsequent refolding of HR2 onto HR1 leads to the formation of the six-helix bundle core structure ^22–24^, which pulls the viral membrane into close apposition to the host-cell membrane and, thus, sets the stage for membrane fusion ^22^.

Membrane fusion generates a lipid intermediate hemifusion state, that is predicted to break and evolve to fusion pore opening ^25^, which is regulated by six-helical bundle formation ^26,27^. Furthermore residues within FPPR, FP, MPER and TM have been as well implicated in fusion ^28–32^ indicating that final steps in fusion are controlled by the conformational transitions of the membrane anchors into the final post-fusion conformation.

Here, we set out to understand the conformational transitions of the gp41 membrane anchors. We show that the presence of the membrane anchors increases thermostability. However, complex formation with a MPER-specific neutralizing nanobody induced an asymmetric conformation of the membrane anchors, which constitutes a late fusion intermediate. We show that this conformation can be targeted by MPER bnAbs consistent with the possibility that MPER-specific nAbs can interfere all along the fusion process until a late stage. Starting from the asymmetric conformation, we used MD simulation based modelling to generate the final post-fusion conformation, which reveals a tight helical interaction of FP and TM in the membrane consistent with its high thermostability. Our work, thus, elucidates the structural transitions of the membrane anchors that are essential for membrane fusion, which can be blocked by MPER-specific bnAbs up to a late stage in fusion.

## Results

### Gp41FP-TM interaction with 2H10

Two gp41 constructs, one containing residues 512 to 581 comprising FP, FPPR and HR1 (N-terminal chain, chain N) and one coding for resides 629 to 715 including HR2, MPER and TM (C-terminal chain, chain C)(**Fig. S1A**) were expressed separately, purified and assembled into the monodisperse trimeric complex gp41FP-TM (**Fig. S1B**). Gp41FP-TM reveals a thermostability of > 95°C as measured by circular dichroism (**Fig. S2A**) indicating that the presence of FP and TMR increases the thermostability by > 7°C compared to gp41 lacking FP and TM ^33^. In order to facilitate crystallization, gp41FP-TM was complexed with the llama nanobody 2H10 ^34^ in β-OG buffer and purified by size exclusion chromatography (SEC)(**Fig. S1C**). To determine the stoichiometry of binding, we performed isothermal titration calorimetry (ITC), which indicated that gp41FP-TM and 2H10 form a 3:1 complex with a K_D_ of 2.1 +/− 0.9 μM (**Fig. S2B**). Interaction of gp41FP-TM with 2H10 was further confirmed by biolayer interferometry (BLI) analysis (**Fig. S2C**).

### Crystal structure of gp41 in complex with 2H10

The structure of gp41FP-TM in complex with 2H10 was solved by molecular replacement to a resolution of 3.8 Å (**Table S1**). The asymmetric unit contained trimeric gp41FP-TM bound to one 2H10 nanobody as indicated by ITC (**Fig. S2B**). The six-helix bundle structure composed of three N-terminal and three C-terminal chains is conserved from HR1 residue A541 to HR2 residue L661 in all three protomers, and identical to previous structures ^22,23^. However, TMR and FP do not follow the three-fold symmetry and their chains point into opposite directions (**Fig. 1A**). 2H10 interacts with chain C-A (**Fig. 1A and B**) and induces a partially extended MPER conformation, including a kink at L669 that positions the rest of MPER and TM (N674 to V693) at a 45° angle with respect to the six-helix threefold symmetry axis. The corresponding N-terminal chain A (chain N-A) has its FP disordered and FPPR from G527 to A533 is flexible, while the remaining FPPR and HR1 form a continuous helix (**Fig. 1C**). The chain C-A 2H10 epitope spans from residues Q658 to N671, which is involved in a series of polar contacts with 2H10. These include interactions of gp41FP-TM E662 to 2H10 Y37, S668 and the carbonyl of D664 to R56, K665 to E95, N671 and the carbonyl of A667 to R54, K655 to R97 and R93 contacts E95 to position it for interaction with K665 (**Fig. 1B**). Notably, mutations of R56, R93, E95 and R97 have been shown to affect interaction ^34^. Chain N-B of the second protomer forms a long continuous helix comprising FP, FPPR and HR1 from residues L518 to D589 with the first six residues of FP being disordered. Likewise, chain C-B folds into a continuous helix from M629 to A700 comprising HR2, MPER and TM (**Fig. 1D**). Cα superimposition of chain C-B with MPER containing gp41 structures ^33,35^ yields root mean-square deviations of 0.55 Å and 0.29 Å (**Fig. S3**), indicating that the straight helical conformation is the preferred conformation in threefold symmetrical gp41. In the third protomer, chain N-C has a helical FP linked by flexible residues G531 to A533 to a short helix of FPPR that bends at A541 with respect to helical HR1. Its corresponding chain C-C contains helical HR2 and a flexible region from N671 to N674, which stabilizes a ~45° rotation of the remaining MPER-TM helix that extends to residue R707 (**Fig. 1E**). Thus, the structure reveals flexible regions within FPPR and MPER. FPPR flexibility is supported by strictly conserved G528 and G531, while MPER has no conserved glycine residues. However, the same kink within L661 to F673 has been observed in the MPER peptide structure ^36^, and in complex with bnAb 10E8 ^37^. The N-terminal FP residues 512 to 517 are disordered within the detergent micelle. Flexibility of this region in the absence of membrane is consistent with NMR peptide structures that propose a flexible coil structure of the N-terminal part of FP in solution followed by a helical region starting at L518 ^38^ as observed here. Based on the flexible linkage of FP and TMR, we propose that both FPPR and MPER act as hinges during gp41 refolding leading to membrane fusion.

**Fig. 1.**
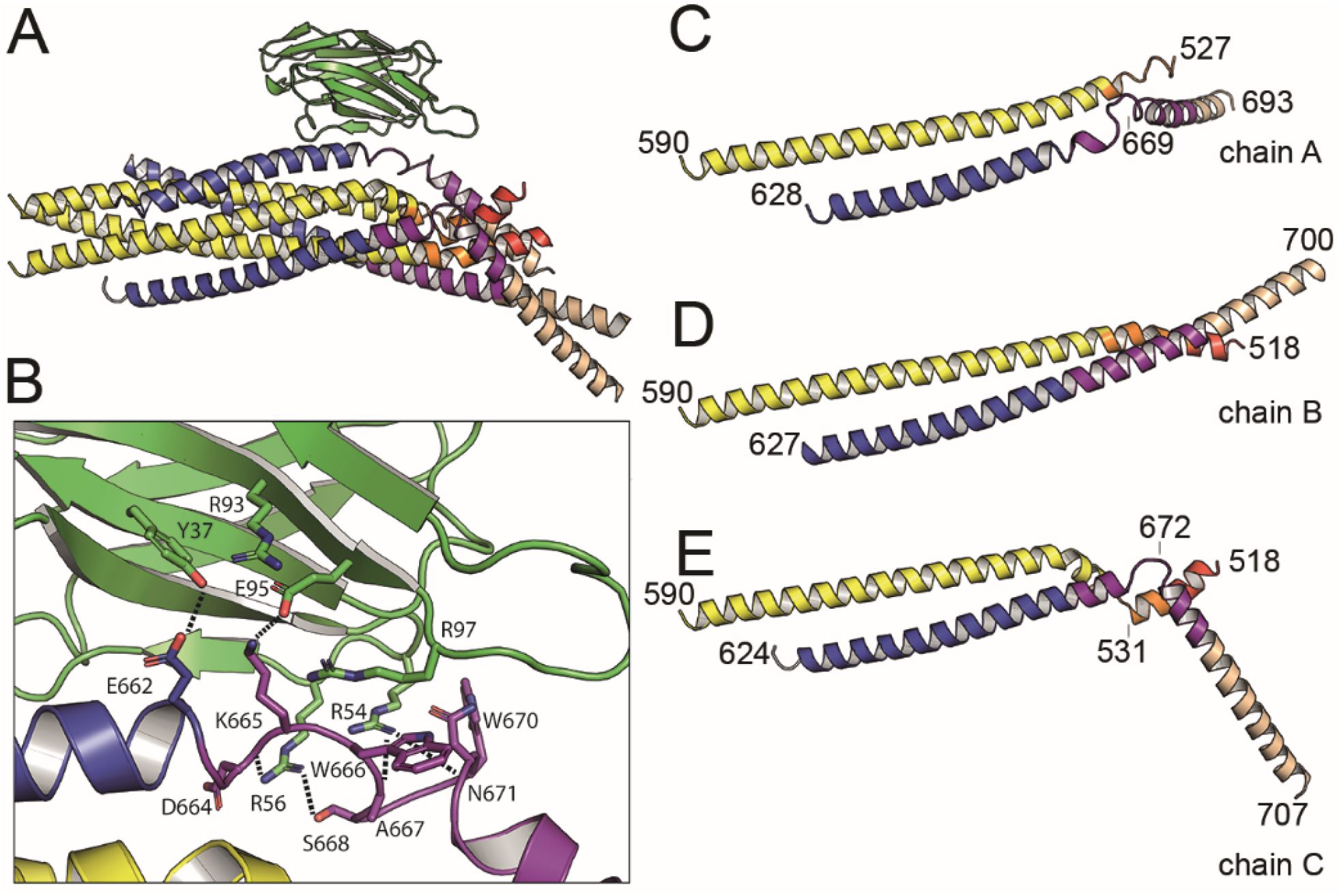
Crystal structure of gp41FP-TM in complex with 2H10. **A,** Ribbon presentation of gp41TM-FP in complex with 2H10. Color-coding of the different segments is as indicated in the gp41 scheme (Fig. S1A), the 2H10 nanobody is colored in green. **B**, Close-up of the interaction of gp41FP-TM with 2H10. Residues in close enough contact to make polar interactions are shown as sticks. **C, D, E,** Ribbon diagram of the individual protomers named chain A, B and C. Residues within the FPPR and MPER hinge regions are indicated.

### MD simulation of gp41FP-TM in a lipid bilayer

In order to test whether the structure is influenced by the presence of the detergent, we probed its stability by MD simulation in a bilayer having the lipid composition of the HIV-1 envelope. This confirmed that the structure is stable in a membrane environment during a 1 μs simulation as only the flexibly linked FP of chain N-C moves within the bilayer during the simulation (**Fig. S4A and B**). The tip of the 2H10 CDR3 dips into the bilayer (**Fig. S4B**), hence confirming the membrane-anchoring role of W100 for neutralization ^34^.

### Neutralization activity of 2H10 depends on membrane interaction

The structure suggests that 2H10 induced the asymmetry within the membrane anchors. Crystal packing effects on the conformation can be excluded, because only the C-terminus of the chain C-C helix is involved in crystal lattice contacts (**Fig. S5**). We therefore further evaluated 2H10 as a neutralizing nanobody, which showed modest neutralization as a bi-head (bi-2H10), whereas neutralization depended on W100 located at the tip of CDR3 ^34^, a hall mark of MPER-specific bnAbs ^39^. In order to improve the breadth and potency of monovalent 2H10, we increased its membrane interaction capacity by changing CDR3 S100d to F (2H10-F) alone and in combination with additional basic residues S27R, S30K and S74R (2H10-RKRF) within the putative 2H10 membrane-binding interface suggested by MD simulation (**Fig. S4C**). Wild type 2H10 did not show significant neutralization against a panel of 10 clade B pseudo-viruses as shown previously ^34^, with the exception of some weak neutralization of NL4-3 and SF163P3. However, both 2H10-F and 2H10-RKRF show improved potency and breadth neutralizing six and eight pseudo-viruses, respectively, albeit with less potency than wild-type bi-2H10 and bnAb 2F5, the latter recognizing an overlapping epitope (**Table 1**). This result, thus, confirms monovalent 2H10 as a modest anti-MPER Ab that neutralizes by engaging MPER and the membrane.

**Table 1.** Pseudovirus neutralization by 2H10, 2H10-F, 2H10-RKRF and bi-2H10. IC50s are indicated in μg/ml.

|  | Tier | 2H10 wt | 2H10-F | 2H10-RKRF | Bi-2H10 | 2F5 | VRC01 |
| --- | --- | --- | --- | --- | --- | --- | --- |
| **NL4-3** | 1 | 25.20 | 18.68 | 9.15 | 1.84 | 0.16 | 0.20 |
| **MN-3** | 1 | >50.00 | 30.38 | 9.36 | 1.39 | 0.03 | 0.06 |
| **BaL.26** | 1 | >50.00 | 19.38 | 9.63 | 6.05 | 1.21 | 0.13 |
| **SF162** | 1a | >50.00 | >50.00 | 25.19 | 6.14 | 1.22 | 0.39 |
| **SF162P3** | 2 | 22.04 | 13.14 | 6.76 | 1.32 | 1.96 | 0.24 |
| **SC422661.8** | 2 | >50.00 | >50.00 | 27.93 | 3.79 | 1.00 | 0.27 |
| **JR-FL** | 2 | 44.65 | 16.93 | 6.95 | 1.49 | 0.97 | 0.11 |
| **JR-CSF** | 2 | >50.00 | 21.66 | 10.85 | 2.85 | 1.24 | 0.37 |
| **QH0692.42** | 2 | >50.00 | >50.00 | >50.00 | >50.00 | 1.20 | 1.21 |
| **THRO4156.18** | 2 | >50.00 | >50.00 | >50.00 | >50.00 | >50.00 | 3.84 |

### 2H10 blocks fusion before the stage of lipid mixing

The efficacy of bi-2H10 and 2H10-RKRF for blocking membrane merging was further assessed in peptide-induced lipid-mixing assays, whereas a vesicle population is primed for fusion by addition of the N-MPER peptide containing the 2H10 epitope, which produces a fluorescence intensity spark at time 20 s (**Fig. 2A**) ^40^. Under these experimental conditions, incorporation of the peptide into the vesicles takes less than 10 sec ^40^. After 120 sec, the mixture was supplemented with target vesicles fluorescently labeled with N-NBD-PE/N-Rh-PE (indicated by the arrow in **Fig. 2A**). The increase in NBD intensity as a function of time followed the mixing of the target vesicle lipids with those of the unlabeled vesicles (kinetic trace labeled ‘+N-MPER’). The increase in NBD fluorescence was not observed when labeled target vesicles were injected in a cuvette containing unlabeled vesicles not primed with peptide (‘no peptide’ trace). Lipid mixing was strongly attenuated when the vesicles primed for fusion with N-MPER were incubated with bi-2H10 before addition of the target vesicles (**Fig. 2A**, +N-MPER/+bi-2H10, dotted trace). Thus, the N-MPER-induced membrane perturbations, which can induce fusion with target membranes, were inhibited by incubation with bi-2H10. Comparison of the kinetics of the lipid-mixing blocking effect of 2H10-RKRF, bi-2H10 and the 2F5 Fab showed that the three antibodies inhibited both the initial rates and final extents of lipid mixing induced by N-MPER (**Fig. 2B**). Using a control MPER peptide lacking the 2H10 and 2F5 epitopes for vesicle priming no inhibition of lipid mixing by 2H10-RKRF, bi-2H10 and 2F5 Fab was observed (**Fig. 2C**), indicating that the inhibitory effects depend on epitope recognition. Fusion inhibition levels estimated as a function of the antibody concentration further confirmed the apparent higher potency exhibited by the bi-2H10 (**Fig. 2D**). Lower concentrations of bi-2H10 compared to 2H10-RKRF were required to attain full blocking of the lipid-mixing process when measured 20 sec (initial rates) or 300 sec (final extents) after target-vesicle injection (**Fig. 2D and E**). The higher inhibitory potency of bi-2H10 indicates an avidity effect, which was also evident when the concentration of the epitope-binding fragments was plotted (**Fig. 2D and E**, empty squares and dotted line). Moreover, bi-2H10 appeared to block efficiently the process even at 2H10:N-MPER ratios below 1:3 (mol:mol), consistent with the involvement of peptide oligomers in the promotion of membrane fusion. Based on these data, we suggest that both 2H10-RKRF and bi-2H10 neutralize HIV-1 at the stage of lipid mixing.

**Fig. 2.**
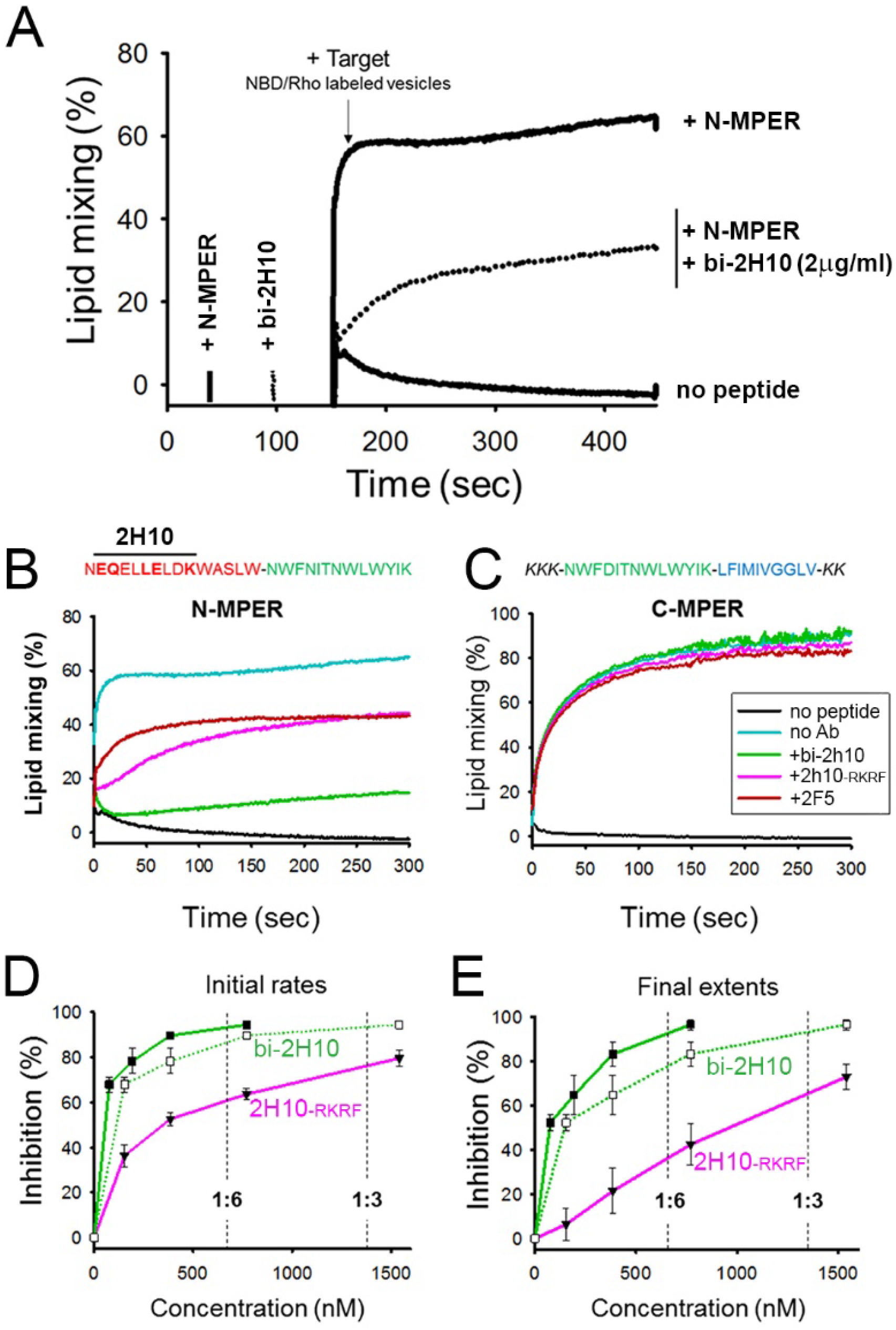
Vesicle-vesicle fusion inhibition by 2H10, bi-2H10 and 2F5. **(A)** Time course of the lipid-mixing assay using fusion-committed vesicles. At time 30 sec (‘+N-MPER’), peptide (4 μM) was added to a stirring solution of unlabeled vesicles (90 μM lipid), and, after 120 sec (indicated by the arrow), the mixture was supplemented with N-NBD-PE/N-Rh-PE-labeled vesicles (10 μM lipid). The increase in NBD fluorescence over time follows the dilution of the probes upon mixing of lipids of target and primed vesicles (+N-MPER trace). NBD increase was substantially diminished in samples incubated with bi-2H10 (2 μg/ml) prior to the addition of the target vesicles (+bi-2H10, dotted trace), and totally absent if unlabeled vesicles were devoid of peptide (‘no peptide’ trace). **(B)** Left: Kinetic traces of N-MPER-induced lipid-mixing comparing the blocking effects of 2H10-RKRF, bi-2H10 and Fab 2F5. **(C)** Absence of effects on lipid-mixing when vesicles were primed for fusion with the C-MPER peptide, devoid of 2H10 and 2F5 epitope sequences. Antibody concentrations were 20 μg/ml in these assays. **(D)** Dose-response plots comparing the inhibitory capacities of 2H10-RKRF and bi-2H10 (purple and green traces, respectively). Levels of lipid-mixing 20 or 300 sec after target vesicle injection were measured (initial rates **D** and final extents, **E**) and percentages of inhibition calculated as a function of the Ab concentration. The dotted line and empty symbols correspond to the effect of bi-2H10 when the concentration of the component 2H10 was plotted. The slashed vertical lines mark the 2H10-to-peptide ratios of 1:6 and 1:3. Plotted values are means±SD of three independent experiments.

### GP41FP-TM interaction with MPER bnAbs

Although the 2H10 epitope overlaps with the 2F5 MPER epitope ^41^, the 2F5-bound peptide structure ^41^, cannot be superimposed without major clashes with adjacent gp41 protomers. In contrast, Cα superposition of the structures of 10E8 and LN01 in complex with MPER peptides demonstrated possible binding to gp41FP-TM chain C-C (**Fig. 3A and B**). Furthermore, HCDR3 of both 10E8 and LN01 could make additional hydrophobic contacts with adjacent FP in this binding mode. To confirm 10E8 and LN01 interaction, we performed immunoprecipitation of gp41FP-TM with both bnAbs, which confirmed their interaction *in vitro* (**Fig. S2D**). We next validated binding by bio-layer interferometry (BLI) using gp41FP-TM as analyte. This revealed KDs of 0,2 nM for 10E8 and 34 nM for LN01 (**Fig. 3C and D**). We conclude that bnAbs 10E8 and LN01 interact with gp41FP-TM with high affinity likely by inducing and stabilizing an asymmetric gp41 conformation similar to the one observed in complex with 2H10 as suggested by the structural modeling (**Fig. 3 and B**).

**Fig. 3.**
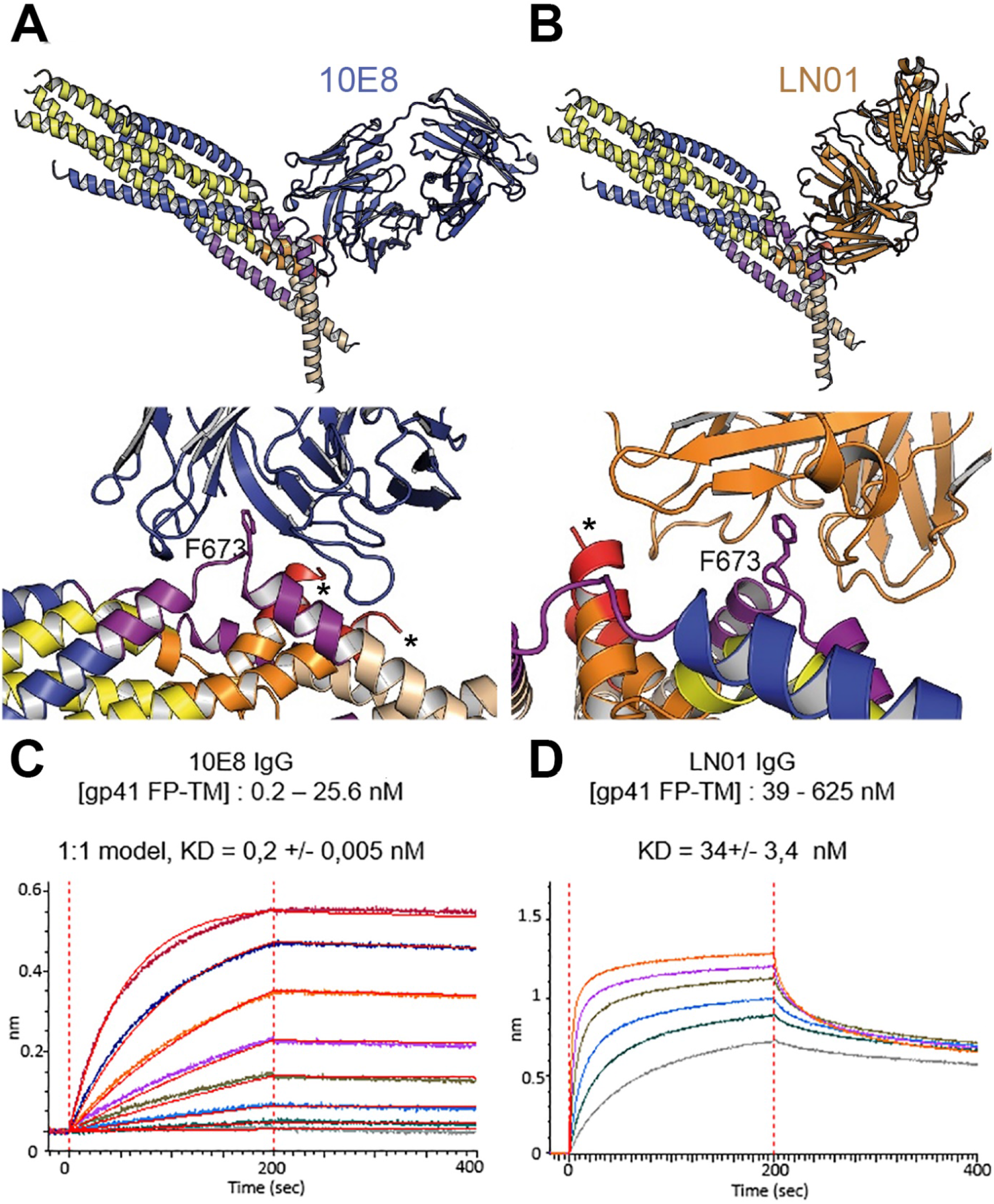
Gp41FP-TM interaction with bnAbs LN01 and 10E8. **A,** Cα superposition of the MPER peptide structure in complex with LN01 (pdb 6snd) onto chain C-C of gp41FP-TM-2H10 structure. The lower panel shows a close-up of the interaction oriented with respect to gp41 F673. **B,** Cα superposition of the MPER peptide structure in complex with 10E8 (pdb 5iq7) onto the corresponding chain C-C of gp41FP-TM. The lower panel shows a close-up of the interaction in the same orientation as in A. **C,** Bio-layer interferometry (BLI) binding of gp41FP-TM to 10E8 and **D,**to LN01. 10E8 binding was fit to 1:1 model and for LN01 a steady state model was employed for fitting the data. For 10E8 binding, gp41FP-TM was used at concentrations from 0.2 to 25,6 nM and for LN01 binding gp41FP-TM concentrations ranged from 39 to 625 nM.

### Building a post fusion conformation by MD simulation

In order to follow the final refolding of the membrane anchors we modeled the post fusion conformation employing MD simulation. Assuming that the final post-fusion conformation shows a straight symmetric rod-like structure we constructed a model of gp41 from the protomer composed of the straight helical chains N-B and C-B (**Fig. S6A and B**). This conformation is also present in the symmetric six-helix bundle structures containing either MPER ^35^ or FPPR and MPER ^33^ (**Fig. S3**). In this model, FP and TM do not interact tightly (**Fig. S6B**), which, however, does not explain the increased thermostability induced by FP and TM (**Fig. S2A**). 1-μs MD simulation of this model (**Fig. S6B**) in solution, rearranges the membrane anchors such that they adopt a compact structure with trimeric FP interacting with adjacent TMs. Furthermore, the TMs kink at the conserved Gly positions 690 and 691, as observed previously ^42^ (**Fig. S6C**). In order to recapitulate the stability of the model in the membrane, we performed an additional 1-μs MD simulation of the model (**Fig. S6C**) in a bilayer resembling the HIV-1 lipid composition, which relaxed the TM to its straight conformation (**Fig. 4A**). The final structural model reveals tight packing of trimeric FP flexibly linked to HR1 by FPPR G525 to G527 (**Fig. 4B**). HR2-MPER and TMR form continuous helices with the TMRs packing against trimeric FP (**Fig. 4A and C**), which spans one monolayer (**Fig. 4A**). As conserved tryptophan residues within MPER have been previously implicated in fusion ^28,30^, we analyzed their structural role in the post fusion model. This reveals that the indole ring of W666 is sandwiched between Leu669 and T536 and packs against L537. W670 makes a coiled-coil interaction with S534, while W672 is partially exposed and packs against L669 and T676. W678 binds into a hydrophobic pocket defined by I675, L679, I682 and adjacent FP/FPPR residues F522 and A526. W680 is partially exposed, but reaches into a pocket created by the flexible FPPR coil (**Fig. S7**). Thus, most of the tryptophan residues have structural roles in the post-fusion conformation, hence providing an explanation for their functional role in fusion ^28^. The MPER epitopes recognized by 10E8 and LN01 are exposed in the post-fusion model, but antibody docking to this conformation produced major clashes, consistent with no expected binding to the final post fusion conformation.

**Fig. 4.**
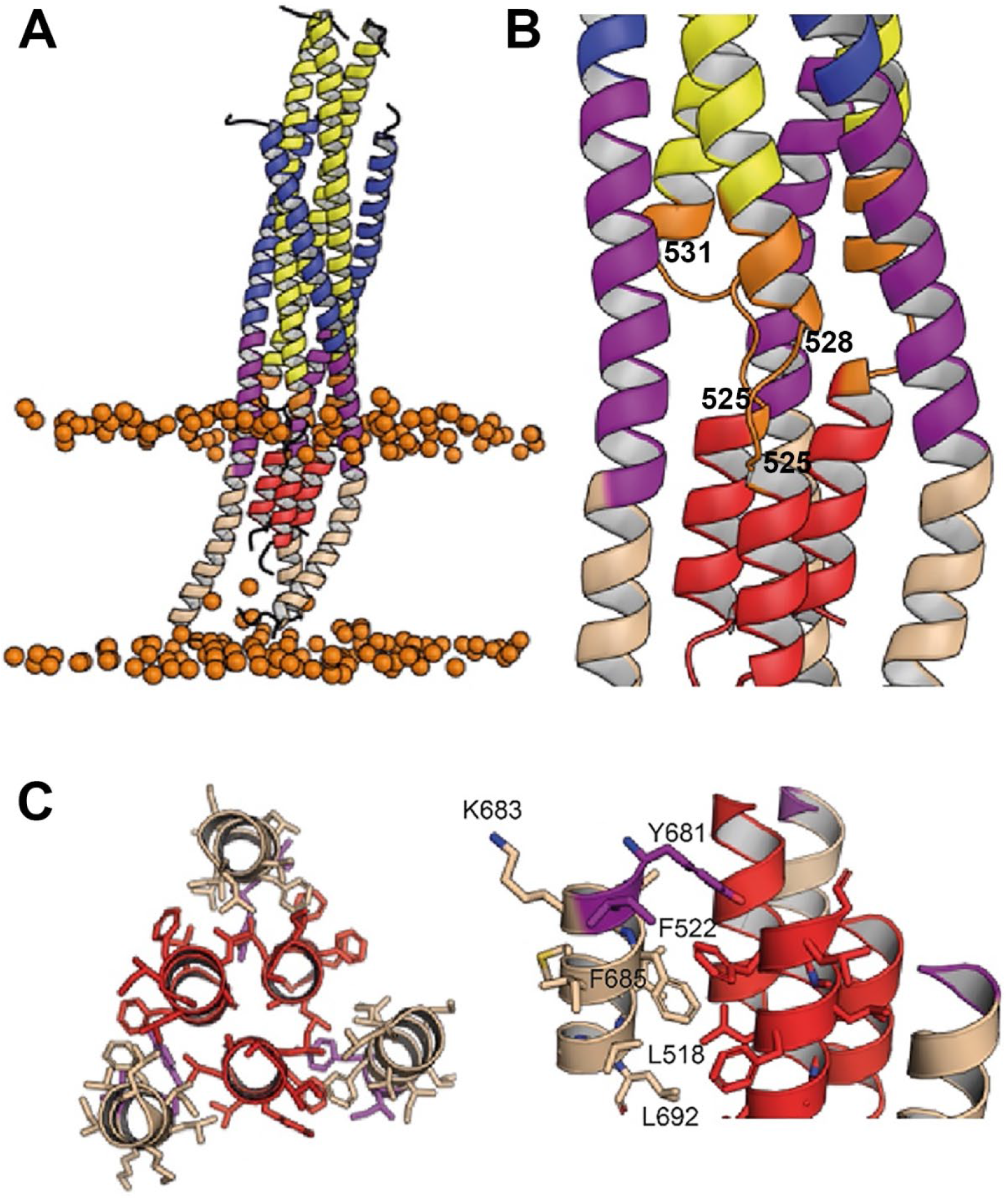
Interactions within the final post fusion conformation of gp41FP-TM modeled by MD. **A**, Model of gp41FP-TM (Fig. S7C) after 1μs MD simulation in a bilayer. Phosphate groups of the phospholipids are shown as orange spheres to delineate the membrane boundaries. **B,** Close up on the MPER and FPPR flexible regions. **C**, Close-up of the interaction of FP (residues 514-524) and TM (residues 681-692) viewed along the three-fold axis from the N-terminus indicating an intricate network of hydrophobic interactions (left panel) and from the side (right panel). Interacting side chains are labeled and shown as sticks.

### Structural transitions of gp41

A number of Env SOSIP structures revealed the native conformation of gp41 (**Fig. 5A and C**) ^9,43,44^. The gp41FP-TM crystal structure and the model of its post fusion conformation provide further insight into the path of conformational changes that native gp41 must undergo to adopt its final lowest energy state conformation. The first major conformational changes in gp41 that take place upon receptor binding are extension of HR1, FPPR and FP into a triple stranded coiled coil with flexible linkers between FPPR and FP that projects FP ~115 Å away from its starting position (**Fig. 5D**). Notably, such an early intermediate fusion conformation structure has been reported for influenza hemagglutinin (HA) ^45^. This is likely followed by an extension and rearrangement of HR2 and MPER producing 11-15-nm long intermediates that connect the viral and cellular membranes ^20,46^. Gp41 refolding into the six-helix bundle structure then produces flexibly linked asymmetric conformations of FPPR-FP and MPER-TMR anchored in the cellular and viral membranes, respectively, as indicated by the gp41FP-TM structure. This intermediate conformation may bring viral and cellular membranes into close proximity (**Fig. 5E**) or may act at the subsequent stage of hemifusion (**Fig. 5F**). Further refolding and interaction of FP-FPPR and MPER-TM will generate the stable post fusion conformation (**Fig. 5G**), a process that completes membrane fusion.

**Fig. 5.**
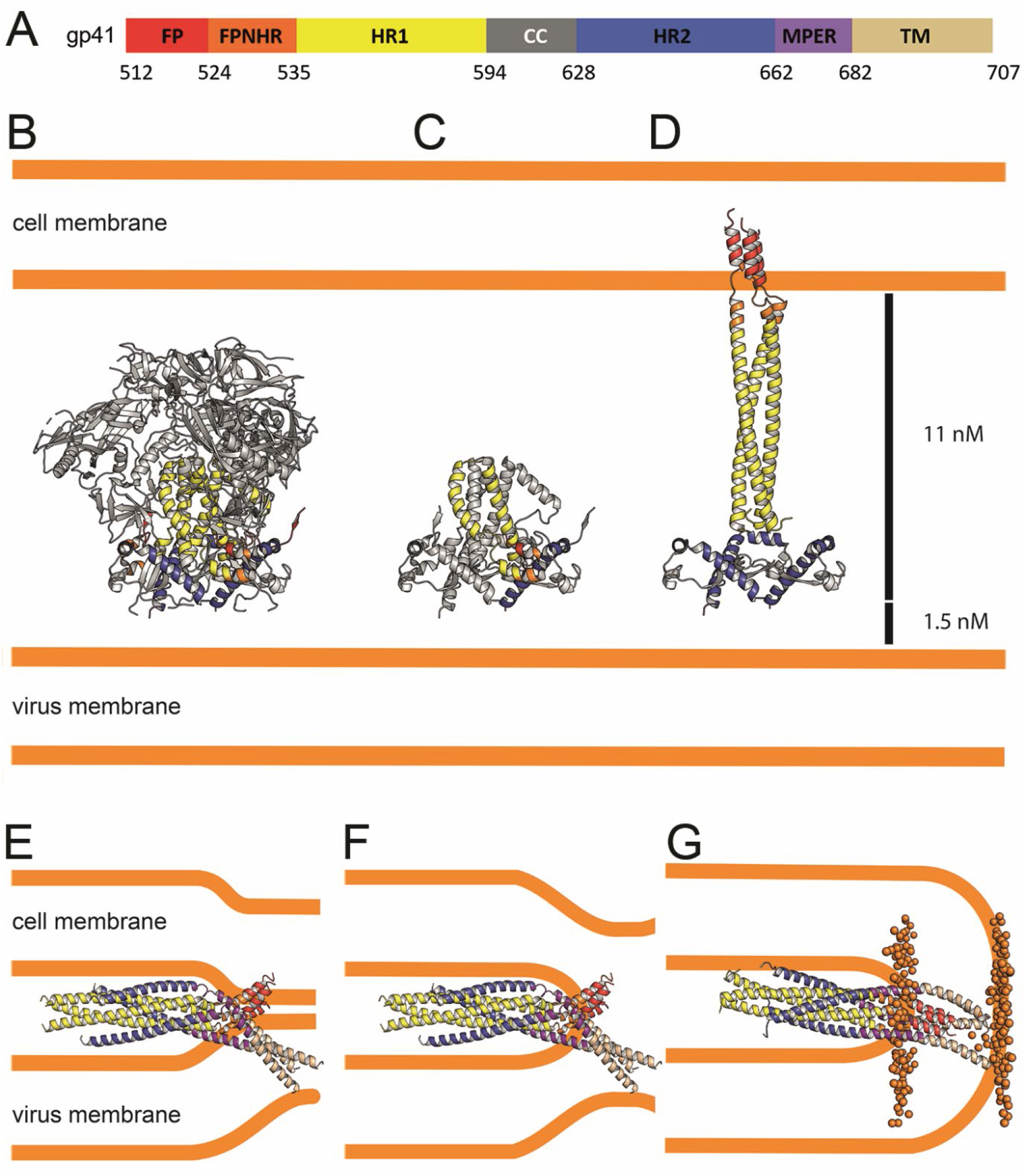
Conformational transitions of gp41 that lead to membrane apposition and membrane fusion. **A,** Representation of the different domains of gp41 with the residue numbers delimiting each domain as indicated. The same color code has been used in all the figures. **B,** Ribbon presentation of the Env prefusion conformation (pdb 5fuu), gp41 is constrained by gp120 in its native conformation. The structure of native gp41 lacks the MPER and TM regions. MPER is spanning a distance of 1.5 nm ^98^. **C**, Ribbon of native gp41, one chain is colored according to the sheme in A and the other two chains are shown in grey. **D,** Binding to cellular receptors CD4 and subsequently to CXCR4/CCR5 induces a series of conformational changes that eventually leads to the dissociation of gp120. During this process, HR1, FPPR and FP will form a long triple stranded coiled coil extending 11 nm towards the target cell membrane. In a first step HR2 may keep its prefusion conformation in analogy to a similar intermediate, activeted influenza virus HA structure ^45^. Alternatively, HR2 may dissociate and form a more extended conformation in agreement with locked gp41 structures bridging viral and cellular membranes that bridge distances of 11 to 15 nm ^46^. **E,** Bending of HR1 and HR2 will result in the six-helical bundle core structure bringing cellular and viral membranes into close apposition with the 3 FPs anchored in the cellular membarne and the 3 TMs anchored in the viral membrane, the gp41 conformatio represented by the gp41FP-TM structure. This intermedaite gp41 conformation may have brought both membranes into close apposition or may have already induced hemifuison as indicated in **F**. **G,** Further reolding of FPPR-FP and MPER-TM results in the final extremely stable post fusion conformation. This thus suggests that rearrangment of the membrane anchors plays crucial roles in lipid mixing, breaking the hemifusion diaphragm to allow fusion pore opening. Boundaries of the lipid layers are shown with orange sphere representing the phosphate atomes of the lipids present in the MD simulation (snapshop taken after 1μs MD simuation).

## Discussion

Membrane fusion is an essential step of infection for enveloped viruses such as HIV-1, and requires extensive conformational rearrangements of the Env prefusion conformation ^7–9^ into the final inactive post-fusion conformation ^2,3^. The fusion model predicts that six-helix bundle formation apposes viral and cellular membranes with FP and TM inserted asymmetrically in the cellular membrane and the viral membrane, respectively ^21^. Here, we show that gp41 containing its membrane anchors can adopt this predicted conformation, which is facilitated by flexible hinges present in FPPR and MPER, thus corroborating their essential roles in membrane fusion ^3,5,47^. The asymmetric conformation of the membrane anchors suggest further that bundle formation occurs before pore formation as suggested previously ^26,27^. The membrane-fusion model proposes further that final steps in fusion involves rearrangement and interaction of TM and FP ^21^, which is confirmed by the MD-simulation model of the post-fusion conformation. Furthermore, the length of the rod-like post-fusion structure of 13 nm lacking the C-C loop is consistent with the gp41 structure lacking FP and TM ^48^.

FP is helical in the gp41FP-TM-2H10 complex and the MD-based post-fusion conformation, in agreement with NMR-based helical FP peptide models ^49,50^, although β-strand structures of FP have been implicated in fusion as well ^51^. In comparison, in native Env conformations, FP adopts multiple dynamic conformations that are recognized by broadly neutralizing antibodies ^43,44,52,53^. In the post-fusion conformation, FP spans one monolayer of the membrane, in contrast to suggested amphipathic helix-like interaction of FP with the outer layer of the target cell membrane ^54,55^.

Furthermore, the coiled-coil interactions within FP and with TM in the post fusion model explain the increased thermostability of gp41FP-TM compared to gp41 lacking FP and TM ^33^. We propose that refolding of FP and TM can liberate additional free energy to catalyze final steps of fusion. Hence, replacement of TM by a phosphatidylinositol (PI) anchor inhibits membrane fusion ^56,57^, akin to the GPI-anchored HA inhibition of influenza virus membrane fusion at the stage of hemifusion ^58^.

Mutations in MPER and FPPR interfere with fusion ^28,30,59,60^, and mutations in TM block fusion ^31^ or reduce fusion efficiency ^32^. Our structural model of the post-fusion conformation predicts that most of these mutations affect the final post-fusion conformation, in agreement with proposed interactions of FPPR and MPER, as well as FP and TM ^61,62^, thereby corroborating their essential roles at late stages of membrane fusion.

Gp41FP-TM interaction with the 2H10 MPER-specific nanobody induces the asymmetric conformation of the membrane anchors. In order to confirm that 2H10 is, indeed, a neutralizing MPER-specific nanobody, we engineered increased 2H10 membrane binding, which improved breadth and potency of 2H10 neutralization, in agreement with enlarged potency by increasing membrane-binding of 10E8 ^63–65^. This result, thus confirmed 2H10 as a modest anti-MPER neutralizing antibody that recognizes both its linear epitope and membrane ^34^. Consistent with its neutralization capacity, 2H10 inhibits membrane fusion at the stage of lipid mixing like 2F5 and other anti-MPER bnAbs ^40,66^. Moreover, gp41FP-TM interacts with MPER bnAbs 10E8 ^37^ and LN01 ^42^ in agreement with docking both structures onto the kinked chain C-C MPER epitope. Notably, the kink in the MPER peptide in complex with 10E8 ^37^ is similar to the chain C-C kink and present in MPER peptide NMR structures ^36,50^. Furthermore, Ala mutations in the kink (671-674) affect cell-cell fusion and lower virus infectivity ^67^ corroborating the physiological relevance of the kinked conformation. We therefore propose that 10E8 and LN01 binding to gp41FP-TM induces similar asymmetry, as observed in the gp41FP-TM-2H10 structure by sampling the dynamic states of the membrane anchors.

Our data, thus, indicate that MPER antibodies can act all along the gp41 refolding pathway from blocking initial conformations of close to native Env ^68–70^ up to a late fusion intermediate state that has already pulled viral and cellular membranes into close apposition. This thus, opens a long temporal window of action for MPER bnAbs consistent with the findings that the half-life of neutralization of MPER bnAbs is up to 30 minutes post virus exposure to target cells ^71,72^. Furthermore, only one Ab per trimer may suffice to block final refolding of the membrane anchors required for fusion. Finally, the presence of dynamic linkers connecting the core of viral fusion proteins with their membrane anchors FP and TM must be a general feature of all viral membrane fusion proteins.

## Materials and Methods

### Cell Lines

TZM-bl cells were obtained from NIH-AIDS Research and Reference Reagent Program (ARRRP) and used for neutralization assays. TZM-bl cells were maintained in Dulbecco’s modified Eagle’s medium supplemented with 10% fetal bovine serum, 100 units of Penicillin and 0.1 mg/ml of Streptomycin while TZM-bl expressing the FcγRI cells were maintained in Dulbecco’s modified Eagle’s medium supplemented with 10% fetal bovine serum, 0.025M Hepes, 50 μg/ml of Gentamicin, 20 μg/ml of Blasticidin.

### HIV-1 Primary Viruses

Env-pseudotyped viruses were prepared by co-transfection of HEK 293-T cells with plasmids encoding the respective *env* genes and the luciferase reporter HIV vector pNLluc-AM as described ^73^. A full list of Env pseudotyped viruses generated with corresponding gene bank entry, subtype and Tier information is provided in Table S2.

### GP41 expression and purification

DNA fragments encoding HIV-1 Env glycoprotein amino acids 512 to 581 (N-terminal chain, chain N) and residues 629 to 715 (C-terminal chain, chain C) were cloned into vectors pETM20 and pETM11 (PEPcore facility-EMBL), respectively. Both constructs contain an N-terminal Flag-tag (DDDDK sequence) and chain C contains additional two C-terminal arginine residues (Fig. S1A). Proteins were expressed separately in *E. coli* strain C41(DE3)(Lucigen). Bacteria were grown at 37°C to an OD_600nm_ of 0,9. Cultures were induced with 1mM IPTG at 37°C for 3h for gp41 chain N and at 25°C for 20h for gp41 chain C. Cells were lysed by sonication in buffer A containing 20 mM Tris pH 8, 100 mM NaCl and 1% CHAPS (3-[(3-cholamidopropyl) diméthylammonio]-1-propanesulfonate (Euromedex). The supernatant was cleared by centrifugation at 53 000 g for 30 min. Gp41 chain N supernatant was loaded on a Ni^2^-sepharose column, washed successively with Buffer A containing 1M NaCl and 1M KCl, then Buffer A containing 50 mM imidazole. Gp41 chain N was eluted in Buffer A containing 500 mM imidazole. Gp41 chain C was purified employing the same protocol as for gp41 chain N. Gp41 chain N was subsequently cleaved with TEV (Tobacco Etch Virus) protease for 2h at 20°C and then overnight at 4°C. After buffer exchange with a mono Q column using buffer B (Buffer A with 0,5 M NaCl), uncleaved material and cleaved His-tags were removed by a second Ni^2+^-sepharose column in buffer A. TEV-cleaved gp41 chain C and chain N were then mixed in a molar ratio 4:1 and incubated overnight. To remove the excess of gp41 chain C, the gp41 complex was loaded on a 3rd Ni^2+^-sepharose column in buffer A, washed with buffer A containing 50 mM imidazole and eluted with buffer A containing 500 mM imidazole. Subsequently the gp41 chain N TrxA-His-tag was removed by TEV digestion for 2h at 20°C and overnight at 4°C. After buffer exchange with a mono Q column in buffer B uncleaved complex and the TrxA-His-tag fusion were removed by a 4th Ni^2+^-sepharose column. The final gp41FP-TM complex was concentrated and loaded onto a Superdex 200 size exclusion column (SEC) in buffer C containing 20 mM Tris pH 8,0, 100 mM NaCl and 1% n-octyl β-D-glucopyranoside (Anatrace).

### Nanobody 2H10 expression

2H10 encoding DNA was cloned into the vector pAX51 ^34^ and expressed in the *E. coli* BL21(DE3) strain (Invitrogen). Bacteria were grown at 37°C to an OD_600nm_ of 0,7 and induced with 1mM IPTG at 20°C for 20h. After harvesting by centrifugation, bacteria were resuspended in lysis buffer containing 20 mM Hepes pH 7,5 and 100 mM NaCl. Bacteria were lysed by sonication and centrifuged at 48 000g for 30 min. Cleared supernatant was loaded on Protein A sepharose column, washed with lysis buffer and eluted with 0,1 M glycine pH 2,9. Eluted fractions were immediately mixed with 1/5 volume of 1M Tris pH 9,0. 2H10 was then further purified by SEC on a superdex 75 column in PBS buffer. Mutants of 2H10, 2H10-F (S100d) and 2H10-RKRF (S27R, S30K, S74R and S100d) were synthesized (Biomatik) and purified as described for the wild type. The 2H10 bi-head was purified as described ^34^.

### Circular dichroism

CD measurements were performed using a JASCO Spectropolarimeter equipped with a thermoelectric temperature controller. Spectra of gp41-TM were recorded at 20 °C in 1 nm steps from 190 to 260 nm in a buffer containing PBS supplemented with 1% n-octyl β-D-glucopyranoside. For thermal denaturation experiments, the ellipticity was recorded at 222 nm with 1°C steps from 20° to 95°C with an increment of 80°C h^−1^, and an averaging time of 30 s/step. For data analysis, raw ellipticity values recorded at 222 nm were converted to mean residue ellipticity.

### Isothermal Titration Calorimetry (ITC)

The stoichiometry and binding constants of 2H10 binding to gp41 FP-TM was measured by ITC200 (MicroCal Inc.). All samples used in the ITC experiments were purified by SEC in a buffer containing 20 mM Tris pH 8.0, 100 mM NaCl and 1 % n-octyl β-D glucopyranoside and used without further concentration. Samples and were equilibrated at 25 °C before the start of the experiment. The ITC measurements were performed at 25 °C by making 20 2-μl injections of 267 μM 2H10 to 0.2 ml of 19.5 μM gp41FP-TM. Curve fitting was performed with MicroCal Origin software. Three experiments were performed, with an average stoichiometry N = 1.1 +/− 0.2 2H10 binds to gp41FP-TM with a KD of 2.1 μM +/− 0.9.

### Bio-layer Interferometry Binding Analysis

Binding measurements between antibodies (10E8 IgG, LN01 IgG and 2H10) were carried out on an Octet Red instrument (ForteBio). For the determination of the binding between antibodies and gp41FP-TM, 10E8 IgG or LN01 IgG or 2H10 were labelled with biotin (EZ-Link NHS-PEG4-Biotin) and bound to Streptavidin (SA) biosensors (ForteBio). The biosensors loaded with the antibodies were equilibrated in the kinetic buffer (20 mM Tris pH 8.0, 100 mM NaCl and 1 % n-octyl β-D glucopyranoside) for 200-500 sec prior to measuring association with different concentrations of gp41FP-for 100-200 seconds at 25 °C. Data were analyzed using the ForteBio analysis software version 11.1.0.25 (ForteBio). For 10E8 the kinetic parameters were calculated using a global fit 1:1 model and 2:1 model. For the determination of the binding of LN01 IgG and 2H10, KDs were estimated by steady state analysis. All bio-layer interferometry experiments were conducted at least three times.

### Immunoprecipitation of gp41FP-TM by bnAbs 10E8 and LN01

220 μg of Gp41FP-TM were incubated alone or with 50μg of 10E8 or LN01 antibodies for 10 h at 20°C in buffer C. The complex was loaded on Protein A sepharose affinity resin and incubated for 1h. The resin was subsequently washed 3 times with buffer C and eluted with SDS gel loading buffer and boiling at 95°C for 5 min. Samples were separated on a 15% SDS-PAGE and stained with Coomassie brilliant blue.

### Neutralization assay

The neutralization activity of the 2H10 variants and mAbs was evaluated using TZM-bl cells and Env pseudotyped viruses as described ^73^. Briefly, serial dilutions of inhibitor were prepared in cell culture medium (DMEM with 10% heat-inactivated FBS, 100 U/ml penicillin and 100 μg/ml streptomycin (all from Gibco)) and added at a 1:1 volume ratio to pseudovirus suspension in 384 well plates (aiming for 500’000–5’000’000 relative light units (RLU) per well in the absence of inhibitors). After one-hour incubation at 37°C, 30 μl of virus-inhibitor mixture was transferred to TZM-bl cells in 384 well plates (6000 cells/well in 30μl cell culture medium supplemented with 20μg/ml DEAE-Dextran seeded the previous day). The plates were further incubated for 48 hours at 37°C before readout of luciferase reporter gene expression on a Perkin Elmer EnVision Multilabel Reader using the Bright-Glo Luciferase Assay System (Promega).

The inhibitor concentration (referring to the mix with cells, virus and inhibitor) causing 50% reduction in luciferase signal with respect to a reference well without inhibitor (inhibitory concentration IC50) was calculated by fitting a non-linear regression curve (variable slope) to data from two independent experiments using Prism (GraphPad Software). If 50% inhibition was not achieved at the highest inhibitor concentration tested, a greater than value was recorded. To control for unspecific effects all inhibitors were tested for activity against MuLV envelope pseudotyped virus.

### Fusion assay

The peptides used in the fusion inhibition experiments, NEQELLELDKWASLW NWFNITNWLWYIK (N-MPER) and *KKK*-NWFDITNWLWYIKLFIMIVGGLV-*KK* (C-MPER), were synthesized in C-terminal carboxamide form by solid-phase methods using Fmoc chemistry, purified by reverse phase HPLC, and characterized by matrix-assisted time-of-flight (MALDI-TOF) mass spectrometry (purity >95%). Peptides were routinely dissolved in dimethylsulfoxide (DMSO, spectroscopy grade) and their concentration determined by the Biscinchoninic Acid microassay (Pierce, Rockford, IL, USA).

Large unilamellar vesicles (LUV) were prepared following the extrusion method of Hope et al. ^74^. 1-palmitoyl-2-oleoylphosphatidylcholine (POPC) and cholesterol (Chol) (Avanti Polar Lipids, Birmingham, AL, USA) were mixed in chloroform at a 2:1 mol:mol ratio and dried under a N2 stream. Traces of organic solvent were removed by vacuum pumping. Subsequently, the dried lipid films were dispersed in 5 mM Hepes and 100 mM NaCl (pH 7.4) buffer, and subjected to 10 freeze-thaw cycles prior to extrusion 10 times through 2 stacked polycarbonate membranes (Nuclepore, Inc., Pleasanton, CA, USA). Lipid mixing with fusion-committed vesicles was monitored based on the resonance energy transfer assay described by Struck et al. ^75^, with the modifications introduced by Apellaniz et al. ^40^. The assay is based on the dilution of co-mixed N-(7-nitro-benz-2-oxa-1,3-diazol-4-yl)phosphatidylethanolamine (N-NBD-PE) and N-(lissamine Rhodamine B sulfonyl)phosphatidylethanolamine (N-Rh-PE) (Molecular Probes, Eugene, OR, USA), whereby dilution due to membrane mixing results in increased N-NBD-PE fluorescence. Vesicles containing 0.6 mol % of each probe (target vesicles) were added at 1:9 ratio to unlabeled vesicles (MPER peptide-primed vesicles). The final lipid concentration in the mixture was 100 μM. The increase in NBD emission upon mixing of target-labeled and primed-unlabeled lipid bilayers was monitored at 530 nm with the excitation wavelength set at 465 nm. A cutoff filter at 515 nm was used between the sample and the emission monochromator to avoid scattering interferences. The fluorescence scale was calibrated such that the zero level corresponded to the initial residual fluorescence of the labeled vesicles and the 100 % value to complete mixing of all the lipids in the system (i.e., the fluorescence intensity of vesicles containing 0.06 mol % of each probe). Fusion inhibition was performed with bi-2H10, 2H10-RKRF and 2F5 Fabs at concentrations of 10 μg/ml and 20 μg/ml as indicated.

### Crystallization, data collection and structure determination

For crystallization, 1 mg of gp41FP-TM was mixed with 1.5 mg of 2H10. The complex was purified by SEC on a Superdex 200 column in a buffer containing 100 mM NaCl, 20 mM Tris pH 8,0 and 1% n-octyl β-D-glucopyranoside and concentrated to 7-10 mg/ml. Crystal screening was performed at the EMBL High Throughput Crystallization Laboratory (HTX lab, Grenoble) in 96-well sitting drop vapor diffusion plates (Greiner). Following manual refinement of crystallization conditions, crystals of gp41FP-TM in complex with 2H10 were grown by mixing 1 μl of protein with 1 μl of reservoir buffer containing 0,1 M sodium citrate pH 6,0, 0,2 M ammonium sulfate, 20% polyethylene glycol 2000 and 0,1 M NaCl at 20°C (293 K) in hanging drop vapor diffusion plates. Before data collection, crystals were flash frozen at 100K in reservoir solution supplemented with 1% n-octyl β-D-glucopyranoside and 25 % ethylene glycol for cryo-protection.

Data were collected on the ESRF beamline ID30b at a wavelength of 0.9730 Å. Data were processed with the program XDS ^76^. The data from two crystals were merged with Aimless ^77^. The data set displayed strong anisotropy in its diffraction limits and was submitted to the STARANISO Web server ^78^. The merged STARANISO protocol produced a best-resolution limit of 3.28 Å and a worst-resolution limit of 4.74 Å at a cutoff level of 1.2 for the local I_mean_ / σ(I_mean_) (note that STARANISO does not employ ellipsoidal truncations coincident with the crystal axes). The gp41FP-TM-2H10 crystals belong to space group C 2 2 2_1_ and the structure was solved by molecular replacement using the program Phaser ^79^ and pdb entries 1env and 4b50. The model was rebuilt using COOT ^80^ and refined using Phenix ^81^. Data up to 3.28 Å were initially used for model building but were finally truncated to 3.8 Å. Statistics for data reduction and structure refinement are presented in Table S1.

One copy of gp41FP-TM in complex with 2H10 are present in the asymmetric unit. Numbering of the nanobody 2H10 was performed according to Kabat. The gp41FP-TM-2H10 complex was refined to 3.8 Å data with an R/Rfree of 26.7 / 31.1 %. 99.6 % of the residues are within the most favored and allowed regions of a Ramachandran plot ^77^. Some of the crystallographic software used were compiled by SBGrid ^82^. Atomic coordinates and structure factors of the reported crystal structures have been deposited in the Protein Data Bank (https://www.rcsb.org; PDB: 7AEJ.

### Figure Generation

Molecular graphics figures were generated with PyMOL (W. Delano; The PyMOL Molecular Graphics System, Version 1.8 Schrödinger, LLC, http://www.pymol.org).

### Molecular Dynamics (MD) simulation

#### Molecular assays

Starting from the crystal structure determined herein, we built two molecular assays based (i) on the structure of the entire Gp41FP-TM/2H10 complex, and (ii) based on a gp41 model generated by a three-fold symmetrization of the straight helical chains N-B and C-B. Electron density for FP and TM is partially absent in the crystal structure and the missing parts have been built as helical extensions; FP from residue 512 to 518 and TM from residues 700 to 709 based on TM structures (6SNE and 6B3U). All residues were taken in their standard protonation state. The first assay included a fully hydrated membrane composed of 190 cholesterol, 40 1-palmitoyl-2-oleoyl-glycero-3-phosphocholine (POPC), 88 1-palmitoyl-2-oleoyl-sn-glycero-3-phospho-ethanolamine (POPE), 36 1-palmitoyl-2-oleoyl-sn-glycero-3-phospho-L-serine (POPS) and 56 N-stearoyl sphingomyelin, present in the HIV-1 lipid envelope ^83^, using the CHARMM-GUI interface ^84,85^. The resulting molecular assembly consisted of about 178,000 atoms in a rhomboidal cell of 106 × 106 × 169 Å³. The second computational assay featured a water bath of 91 × 91 × 114 Å³, representing a total of nearly 95,700 atoms. Both assays were electrically neutral, with a NaCl concentration set to 150 mM.

#### Molecular Dynamics

All simulations were performed using the NAMD 2.14 program ^86^. Proteins, cholesterol, lipids and ions were described using the CHARMM forcefield ^87–89^ and the TIP3P model ^90^ was used for water. MD trajectories were generated in the isobaric-isothermal ensemble at a temperature of 300 K and a pressure of 1 atm. Pressure and temperature were kept constant using the Langevin thermostat and the Langevin piston method ^91^, respectively. Long-range electrostatic interactions were evaluated by the particle-mesh Ewald (PME) algorithm ^92^. Hydrogen mass repartitioning ^93^ was employed for all simulations, allowing for using a time step of 4 fs. Integration was performed with a time step of 8 and 4 fs for long- and short-range interactions, respectively, employing the r-RESPA multiple time-stepping algorithm ^94^. The SHAKE/RATTLE ^95,96^ was used to constrain covalent bonds involving hydrogen atoms to their experimental lengths, and the SETTLE algorithm ^97^ was utilized for water.

The computational assay formed by gp41 in an aqueous environment was simulated for a period of 1 μs, following a thermalization of 40 ns. For the gp41FP-TM/2H10 complex, the lipid bilayer was first thermalized during 200 ns using soft harmonic restraints on every dihedral angle of the protein backbones, allowing the complex to align optimally with its membrane environment. Following the equilibration step, a production run of 1 μs was performed.

The final structure of the hydrated gp41 was also embedded in the HIV-1-like envelope membrane employed for the gp41FP-TM/2H10 complex. The same 200 ns equilibration protocol was applied followed by a production run of 1 μs.

## Acknowledgement

W.W. acknowledges support from the Institute Universitaire de France (IUF), from the European Union’s Horizon 2020 research and innovation programme under grant agreement No. 681137, H2020 EAVI and the platforms of the Grenoble Instruct-ERIC center (IBS and ISBG; UMS 3518 CNRS-CEA-UGA-EMBL) within the Grenoble Partnership for Structural Biology (PSB). Platform access was supported by FRISBI (ANR-10-INBS-05-02) and GRAL, a project of the University Grenoble Alpes graduate school (Ecoles Universitaires de Recherche) CBH-EUR-GS (ANR-17-EURE-0003). J.L.N. acknowledges funding from Spanish MINECO (BIO2015-64421-R; MINECO/AEI/FEDER, UE), Spanish MCIU (RTI2018-095624-B-C21; MCIU/AEI/FEDER, UE) and Basque Government (IT1196-19). We thank Miriam Hock and Serafima Guseva for previous contributions to the project, the ESRF-EMBL Joint Structural Biology Group for access and support at the ESRF beam lines, J. Marquez (EMBL) from the HTX crystallization facility, C. Mas and J.-B. Reiser for assistance on ISBG platforms.

## Author Contributions

W.W. conceived and designed the study. J.L.N. supervised the lipid mixing experiments performed by J.T. A.T. supervised neutralization assays performed by N.F. C.C. and F.D. performed molecular dynamics simulation experiments. D.G. produced proteins, mutants and performed crystallization and pull-down experiments. C.Ca. performed the structural studies and interaction studies. W.W. wrote the manuscript with input from all authors.

