## Supplemental information for "Structure of HIV-1 gp41 with its membrane anchors targeted by neutralizing antibodies"

**Table S1: Crystallographic data collection and refinement statistics.**

| **Data collection** | Gp41FP-TM* |
|---|---|
| Space group | $C\ 2\ 2\ 2_1$ |
| Cell dimensions | |
| *a, b, c* (Å) | 96.75, 101.41, 234.42 |
| α, β, γ (°) | 90, 90, 90 |
| Resolution (Å) | 48.38 - 3.8 (3.94 - 3.8) [1] |
| Unique reflexions | 11179 (631) [1] |
| $R_{merge}$[2] | 0.23 (1.508) [1] |
| $R_{p.i.m}$[3] | 0.081 (0.548) |
| $I / \sigma I$ | 4.75 (1.74) [1] |
| Completeness (%) | 78.01 (54.69) [1] |
| Multiplicity | 9.1 (9.6) [1] |
| CC (1/2) | 0.992 (0.628) [1] |
| **Refinement** | |
| Resolution (Å) | 48.38 - 3.8 (3.936 - 3.8) [1] |
| No. reflections | 9154 (630) [1] |
| Reflections used for $R_{free}$[4] | 550 (51) [1] |
| $R_{work}$[4] / $R_{free}$[5] | 0.267 / 0.312 |
| No. atoms | |
| Protein | 4440 |
| Ligand/ion | 0 |
| Water | 0 |
| Wilson B (Å$^2$) | 82.61 |
| Average B-factors (Å$^2$) | |
| Overall | 91.76 |
| Protein | 91.76 |
| Ligand/ion | |
| Water | |
| R.m.s deviations | |
| Bond lengths (Å) | 0.003 |
| Bond angles (°) | 0.66 |
| Ramachandran Plot (%) | |
| Favored | 96.65 |
| Outliers | 0.37 |
| PDB ID | 7AEJ |

* Data collected from 2 crystals were used for structure determination.
The statistics are for data that were truncated by STARANISO to remove poorly measured reflections affected by anisotropy. $R_{merge}$, $R_{p.i.m}$ and multiplicity are calculated on unmerged data prior to STARANISO truncation. For comparison, after STARANISO truncation, $R_{merge}$ in the resolution shell 3.97 Å - 3.85 Å is 0.787.

[1] Parentheses refer to outer shell statistics.

[2] $R_{merge} = \Sigma_{hkl} \Sigma_i \mid I_{hkl,i} - \langle I_{hkl} \rangle \mid / \Sigma_{hkl} \Sigma_i I_{hkl,i}$, where $I_{hkl,i}$ is the scaled intensity of the ith measurement of

reflection h, k, l, and < Ihkl> is the average intensity for that reflection.
[3] $R_{p.i.m.} = \Sigma_{hkl} \sqrt{1/(n-1)}\, \Sigma_i | I_{hkl,i} - \langle I_{hkl} \rangle | / \Sigma_{hkl} \Sigma_i I_{hkl,i}$,
[4] $R_{work} = \Sigma_{hkl} | Fo - Fc | / \Sigma_{hkl} | Fo | \times 100$, where Fo and Fc are the observed and calculated structures factors.
[5] $R_{free}$ was calculated as for $R_{work}$, but on a test set of 5% of the data excluded from refinement.

**Table S2.** Env pseudoviruses

| HIV-1 Envelope | HIV subtype | Neutralization Tier | Accession Number |
|---|---|---|---|
| NL4_3 | B | 1 | U26942 |
| MN-3 | B | 1 | AY669737 |
| BaL.26 | B | 1 | DQ318211 |
| SF162.LS | B | 1a | EU123924 |
| SF162P3_cl2-4 | B | 2 | AY988107 |
| JR-FL | B | 2 | AY669728 |
| JR-CSF | B | 2 | AY669726 |
| QH0692.42 | B | 2 | AY835439 |
| THRO4156.18 | B | 2 | AY835448 |
| SC422661.8 | B | 2 | AY835441 |

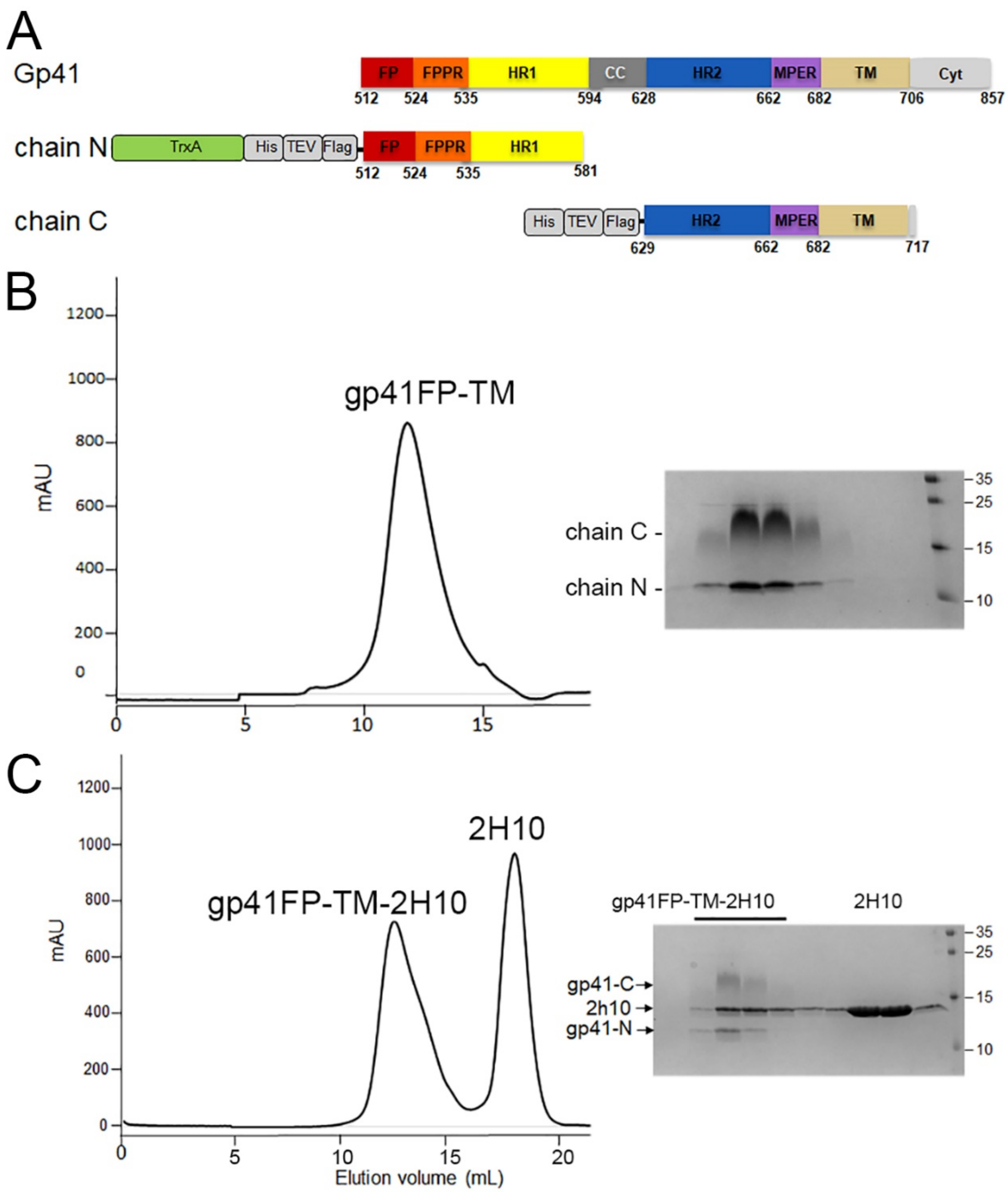

**Fig. S1**. **Characterization of gp41 containing FP and TM.**

**A,** Schematic drawing of gp41 and expression constructs of gp41 chain N and gp41 chain C. Sequence numbering is based on the HIV-1-HBX2 envelope gp160 sequence; TrxA, thioredoxin fusion protein; His, His-tag; TEV, TEV protease cleavage sequence; Flag, flag-tag.

**B**, Size exclusion chromatography of the gp41FP-TM complex composed of chains N and C and SDS-PAGE showing the two bands corresponding to gp41 chains N and C.

**C**, SEC of gp41FP-TM in complex with the llama nanobody 2H10 and corresponding SDS PAGE showing the three bands corresponding to gp41 chains N and C and 2H10.

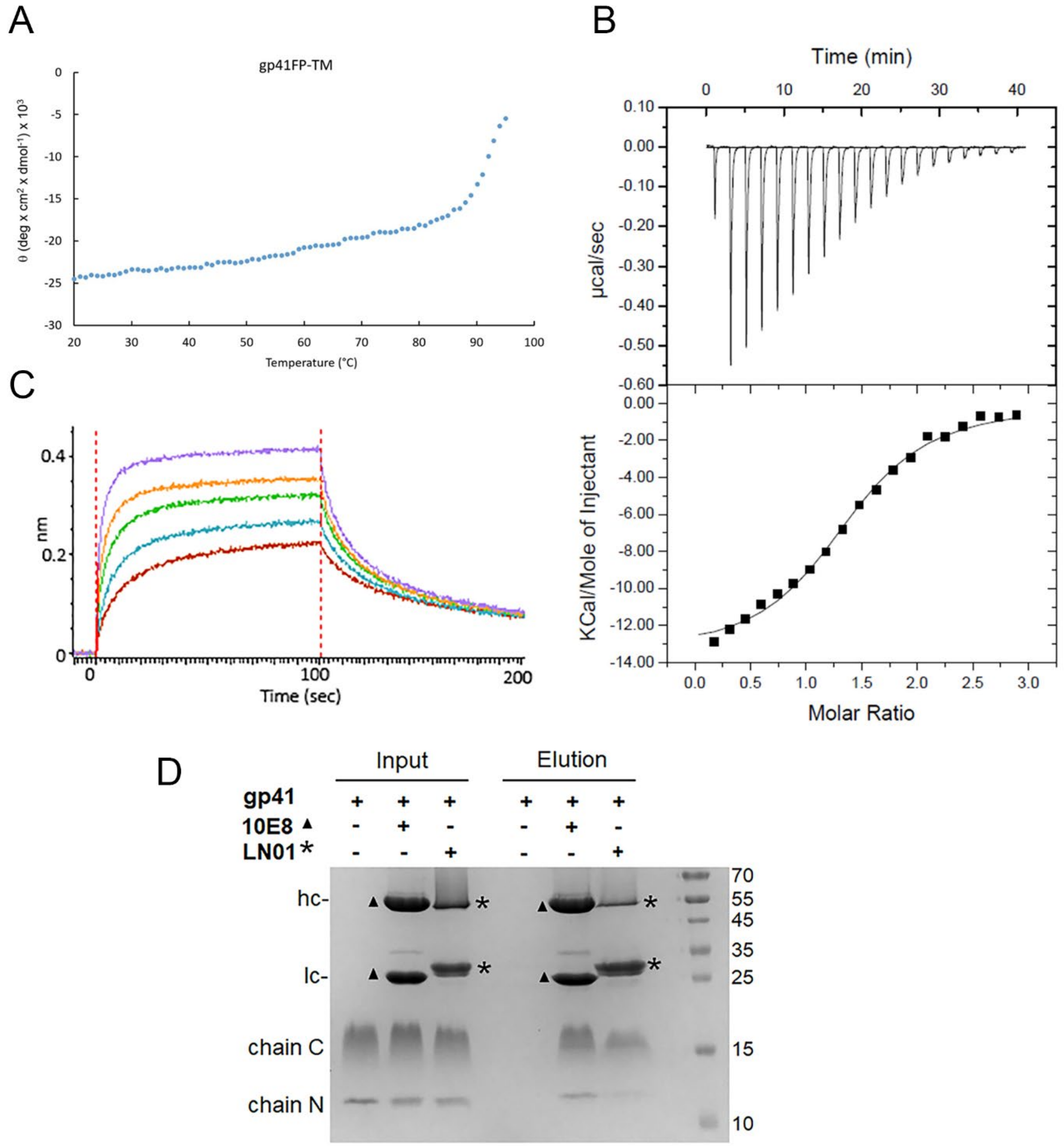

**Fig. S2. Biophysical characterization of gp41FP-TM and MPER Ab interaction**

**A,** Circular dichroism of gp41FP-TM shows that FP and TM increase the melting temperature of gp41. Temperature dependent unfolding of gp41FP-TM monitored by circular dichroism spectroscopy recorded at 222 nm in a buffer containing 1% β-OG. Gp41FP-TM has an estimated Tm of ~95°C.

**B,** Gp41FP-TM complex formation with 2H10. ITC data were recorded on successive injections of 2H10 at a concentration of 267 μM into the cell containing gp41FP-TM at a concentration of 19,5 μM. Three experiments were performed, with an average stoichiometry N = 1.1 +/- 0.2, which suggests that on average only one 2H10 binds to trimeric gp41FP-TM under these conditions. The calculated KD is 2.1 µM +/- 0.9.

**C,** Gp41FP-TM complex formation with 2H10. Bio-layer interferometry (BLI) binding of gp41FP-TM to 2H10. GP41FP-TM concentrations analyzed are dilutions between 156 and 2500 nM. The estimated KD based on the steady state binding model is 170 +/-17 nM. Note that te calculated Kds of the ITC and BLI experiments are only estimates since 2H10 needs to induce the gp41FP-TM binding conformation. Thus, likely, only a fraction of gp41FP-TM may adopt the required conformation during the injection time used to record binding.

**D.** Pull down of gp41FP-TM by bnAb 10E8 and LN01. Immunoprecipitation of gp41FP-TM by bNAbs 10E8 and LN01. Input and eluted fractions were analyzed on SDS-gel and stained with Coomassie brilliant blue. Input fractions correspond to gp41FP-TM alone (lane 1), with 10E8 (lane 2) and with LN01 (lane 3) before incubation with protein A sepharose resin. In absence of antibody, gp41FP-TM is not retained by protein A sepharose (lane 4) but complexes of gp41FP-TM-10E8 and gp41FP-TM-LN01 are eluted from protein A sepharose (lanes 5 and 6, respectively). Bands corresponding to the heavy (hc) and light (lc) chains of 10E8 chains are indicated by ▲ and those of LN01 by ∗. The N- and C-terminal chains of gp41FP-TM are indicated. Molecular weight markers are in kDa.

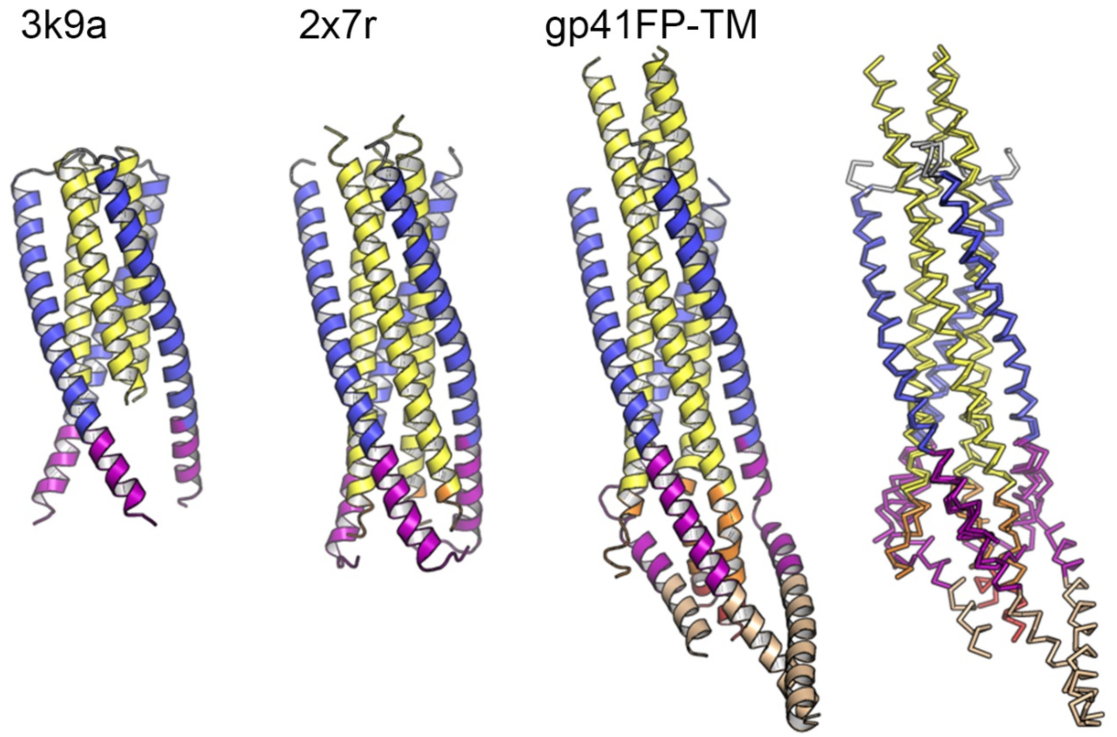

**Fig. S3. Comparison of the gp41FP-TM structure with gp41 core structures.**

Ribbon presentation of gp41-MPER (pdb 3k9a), gp41FPPR-MPER (pdb 2x7r) and gp41FP-TM. Cα super positioning of all three structures onto chains N-B (residues 546 - 574) and C-B (residues 628 - 662) of gp41FP-TM, revealing an r.m.s.d of 0.55 Å between pdb 3k9a and gp41FP-TM and an r.m.s.d. of 0.29 Å between pdb 2x7r and gp41FP-TM for the straight helices of chain B.

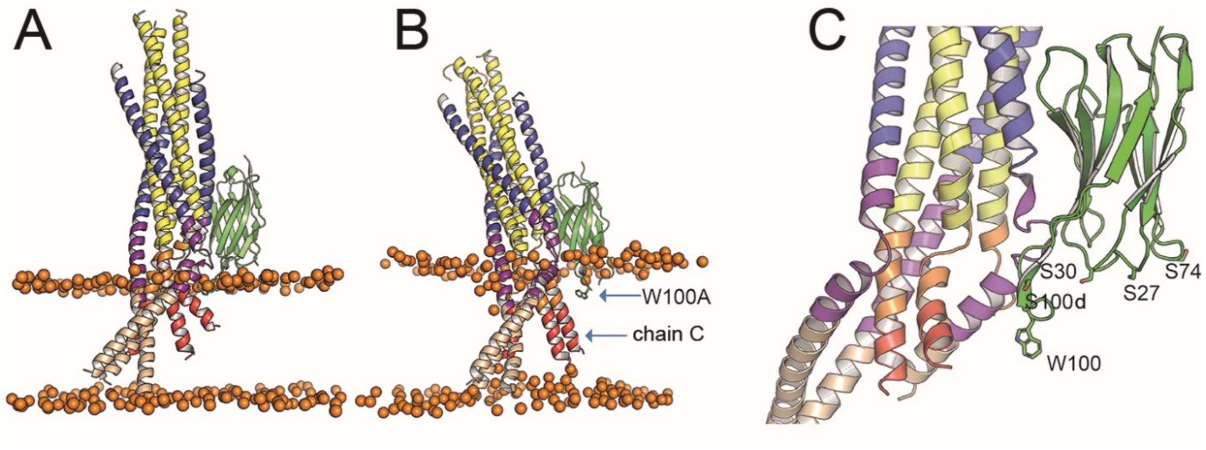

**Fig. S4. Positioning of gp41FP-TM-2H10 in a bilayer by MD simulation.**

**A,** Model of gp41FP-TM-2H10 before simulation and **C**) after 1µs simulation, which repositions the 2H10 CDR3 in the membrane and reveals movement of FP of chain C. The orange spheres represent the phosphate atoms of the phospholipids and mark the membrane boundaries.

**C,** Close-up of the propose membrane interaction of the 2H10 interface. CDR3 W100 and S100d mutated to F could as well insert into the membrane. Furthermore, basic residues at positions S30R, S27R and S74R (shown as sticks) are positioned to make polar interactions with lipid head groups leading to increased membrane binding and thus improved neutralization (2H10-RKRF).

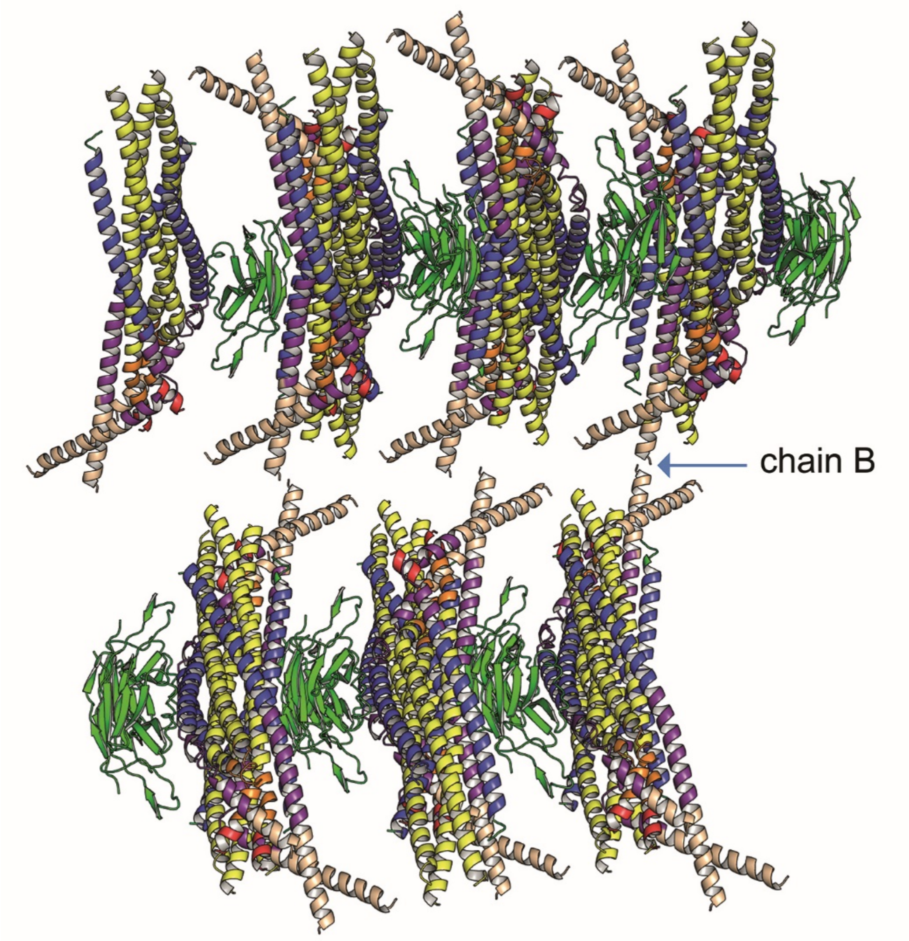

**Fig. S5. Crystal lattice packing.**

Only the TM of chain B makes week crystal contacts between crystal lattices, indicating that the observed asymmetric conformation is not influenced by crystal packing. Note that lateral crystal packing is induced by 2H10 interactions between complexes.

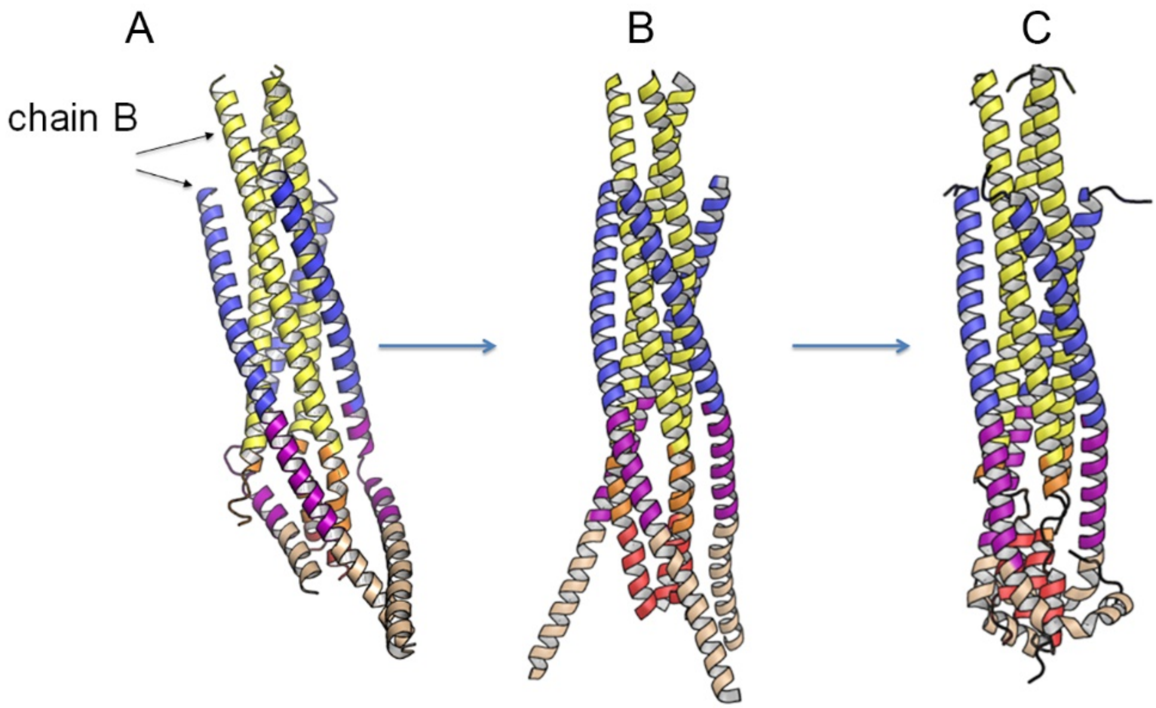

**Fig. S6. Modelling a post fusion conformation by MD simulation.**

**A,** Ribbon of gp41FP-TM.

**B,** Ribbon of the symmetric trimer model built from chains N-B and C-B of the gp41FP-TM structure.

**C,** 1µs MD simulation of the model shown in B, which refolds FPPR-FP and MPER-TM. The kinks in the TM at conserved Gly positions have been observed before {Pinto, 2019 #2827}.

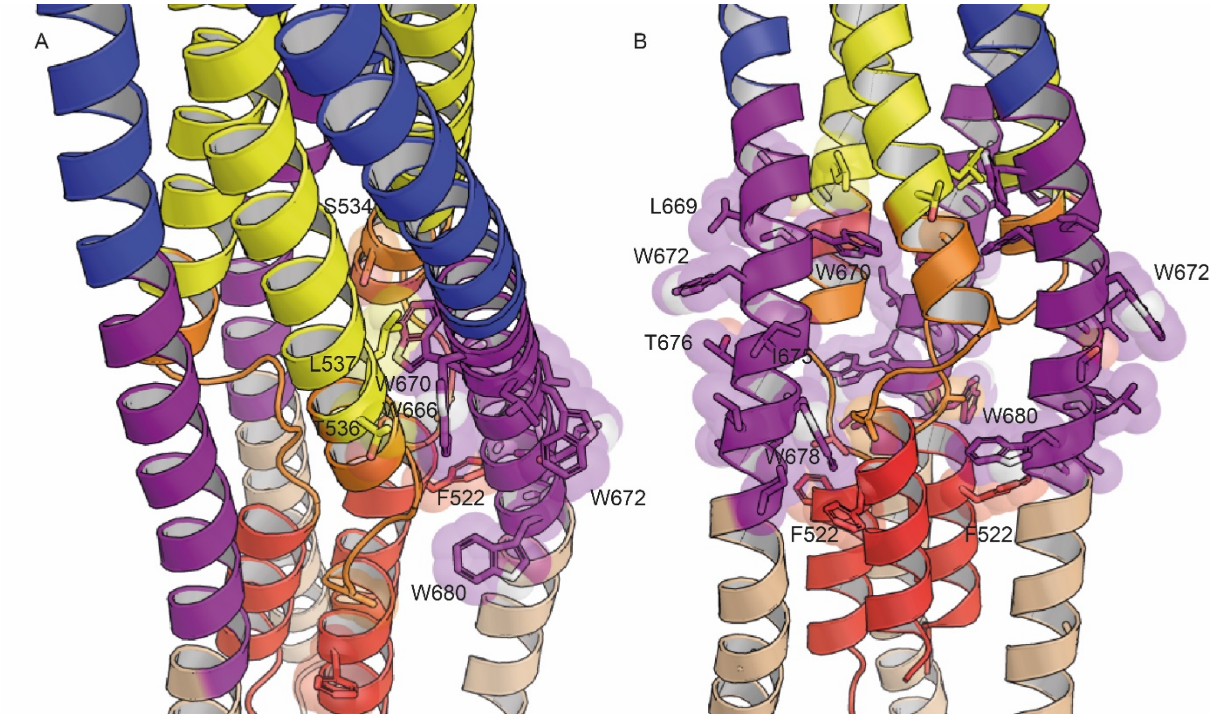

**Fig. S7. Positions of the conserved tryptophan residues of MPER in the post fusion model.** Tryptophan residues W666, W670, W672, W678 and W680 and their close-by potential contacts are shown as spheres.
